# Closed-loop heart rate regulation based on medulla stimulation

**DOI:** 10.64898/2025.11.29.691268

**Authors:** Yizhou Xu, Michele Chan, Masabumi Minami, Yuichi Takeuchi

## Abstract

Cardiovasular diseases are often accompanied with the dysfunctions of autonomic nervous system. Cardiac neuromodulation including vagus nerve stimulation have emerged as alternative therapies by restoring sympatho-vagal balance. In this work, a medulla stimulation strategy was developed to precisely regulate the heart rate. The stimulation parameters effecting on cardiac function were assessed in open-loop experiments. The cardiac effects of rostral and caudal medulla stimulation were enhanced with increased stimulus frequency. We further investigated whether medulla stimulation can temporarily regulate the heart rate at predetermined levels using on-off closed-loop control. Additionally, we tested the capacity of medullary stimulation to rebalance autonomic function under peripheral imbalance conditions caused by dobutamine and neostigmine administration. The performance of heart rate regulation was quantified by the control degree and root mean squared error. The results demonstrate the feasibility of medulla stimulation for regulating sympathetic activity, emphasizing its potential therapeutic use in cardiac disorders.

## Introduction

The autonomic nervous system (ANS) maintains the whole-body physiological function and homeostasis through dynamic interplay between its sympathetic and parasympathetic branches. In the lower brainstem, neurons in the rostral ventrolateral medulla (RVLM) send major excitatory projections to preganglionic sympathetic neurons located in the intermediolateral cell columns in the spinal cord. The vagus nerve originating from the nucleus ambiguous and the dorsal motor nucleus of the vagus nerve provides efferent parasympathetic activity to organs. Dysregulation of sympathetic and parasympathetic tones is associated with development and progression of many cardiovascular diseases.

Pharmacological therapies like beta blockers or ion channel blockers have been the major treatments for cardiovascular disorders. Neuromodulation methods could provide more precisely and timely intervention as alternative treatment for drug-resistant patients^1^. To date, surgery and device-based methods have been developed and applied to modulate ANS activity for cardiovascular diseases as alternative treatments. Carotid baroreceptor stimulation, enhancing baroreflex and exhibiting sympathoinhibition effects, has been shown to be effective in the long-term reduction of blood pressure in resistant hypertension^2–5^. Pulmonary vein isolation has been widely used to prevent depolarizations from the pulmonary vein as a treatment for atrial fibrillation (AF)^6^ and can be combined with renal sympathetic denervation^7,8^ and ganglionated plexi ablation^9,10^. Moreover, renal denervation has been applied to resistant hypertension by regulating kidney natriuresis and the renin-angiotensin system^11–15^. Cervical vagus nerve stimulation has been developed for implantation therapy within bioelectronics medicine. In clinical trials, the vagus nerve stimulation via an implanted device has shown potential applications in the treatment of heart failure^16–20^.

The integration of closed-loop systems in cardiac neuromodulation provided real-time adjustment of stimulation based on a continuous stream of electrocardiogram to regulate the heart rate^21–24^. Closed-loop systems allow for the precise delivery of electrical stimuli when detecting the onset of specific electrical activity in the heart. This approach also provided adaptive therapies for patient-specific needs. This precise and adaptive intervention can significantly improve patient outcomes and minimizes the potential side effects. Previous studies mainly focused on reducing cardiac output through enhanced parasympathetic activity. Whereas precise sympathetic modulation can provide versatile function for future needs, applying to enhance sympathetic activity as treatments or neuroprosthesis. The sympathetic activity was activated under stimulation of the RVLM^25^ and was inhibited by caudal ventrolateral medulla (CVLM) neurons through the baroreflex^26–29^. Taken together, these two regions provide potential targets for sympathetic neuromodulation.

In this study, we explored and evaluated the control of sympathetic activity in medullary stimulation using heart rate as an physiological index. We first assessed the cardiac response by open-loop approach with different stimulation parameters. The closed-loop approach with on-off control method was then applied to temporarily regulate the heart rate (Fig. 1). Finally, the capacity of rebalancing autonomic dysfunction was evaluated following neostigmine and dobutamine injection.

**Figure 1.**
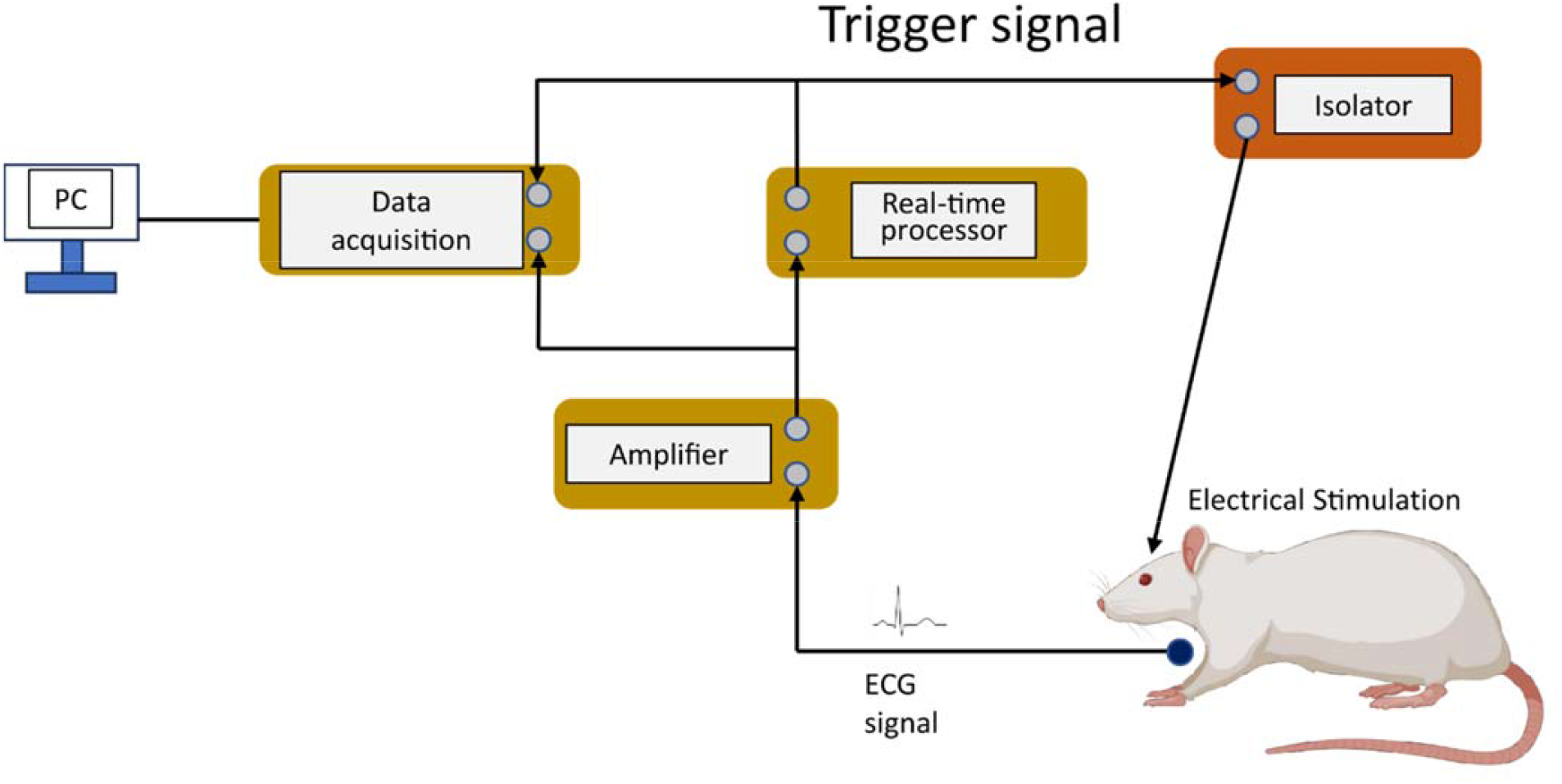
The schematic of the closed-loop stimulation system. Created in BioRender.https://BioRender.com/g7181zt

## Results

### Cardiovascular response in open-loop stimulation

The chronotropic response to medullary stimulation depended on both frequency and intensity, Figure 2 illustrates representative responses to electrical stimulation of the RVLM and CVLM region. Electrical stimulation was delivered with a bipolar 500 μs pulse width. Low intensity stimulation (25 μA) evoked almost no change in basal chronotropic function. Increasing the intensity to 40 μA resulted in tachycardia and bradycardia during the on-phase in the RVLM and CVLM, respectively. Analogously, only high-frequency stimulation induced chronotropic responses during the stimulation. Average chronotropic responses (percentage change from baseline) are summarized in Fig. 2d,e. Chronotropic responses occurred preferentially at higher frequencies, with high frequency providing greater control range and increased sensitivity to stimulus intensity. Additionally, the amplitude of heart rate changes, characterized by the standard deviation of RR interval (SDRR) is shown in Fig. 3a,b. At the high frequency (100 Hz), the stimulation evoked the chronotropic response with a higher peak value, indicating a sustained heart rate change rather than a threshold-like trajectory during the on-phase of stimulation. However, in rostral medulla stimulation using a low frequency of 60 Hz with a high intensity (60 μA), the increased SDRR revealed a slight chronotropic effect, indistinguishable in average percentage change (Fig. 3a). Figure 3d-i also illustrates the change rate of chronotropic response during stimulation using SD1 and SD2 from quantitative Poincaré plots. SD1, representing the variation of adjacent heartbeat, showed no significant difference at all combinations of parameters in both rostral and caudal medulla stimulations. For the long-term variation of SD2, characteristics analogous to SDRR were observed with high frequency, a rapidly induced chronotropic response, and a slight response was demonstrated during stimulation using 60 Hz and 60 μA.

**Figure 2.**
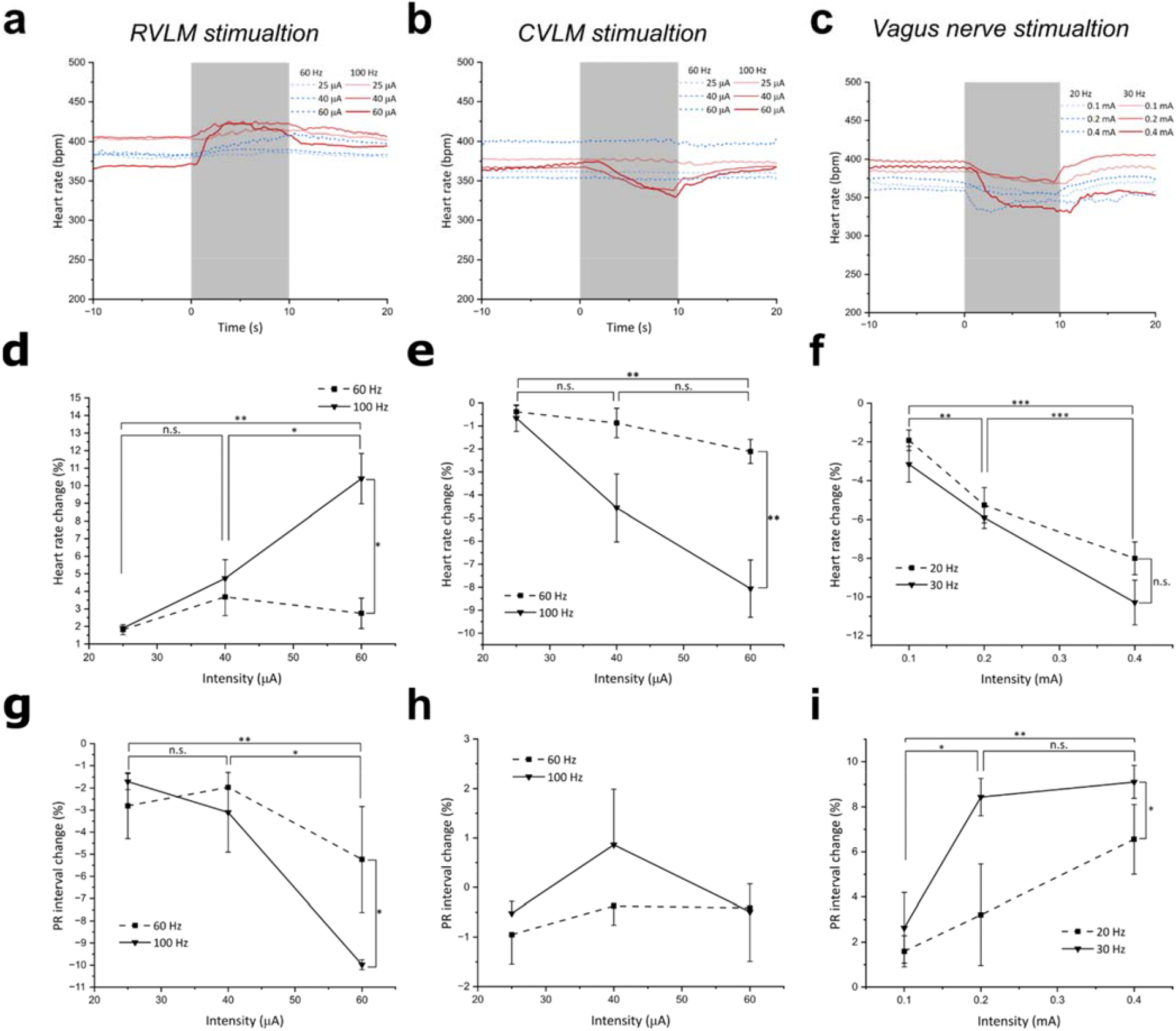
Cardiac effects of open-loop stimulation. Left column, RVLM stimulation; middle column, CVLM stimulation; right column, vagus nerve stimulation. (**a-c**) Typical heart rate trajectory during the electrical stimulation (grey area). (**d-f**) Mean heart rate change from baseline heart rate in electrical stimulation. (**g-i**) Percent changes of PR interval during electrical stimulation. \**P* < 0.05, \*\**P* < 0.01, \*\*\**P* < 0.001, n.s.: not significant by Fisher LSD multiple comparisons test following two-way ANOVA. Data are presented as mean ± SEM.

**Figure 3.**
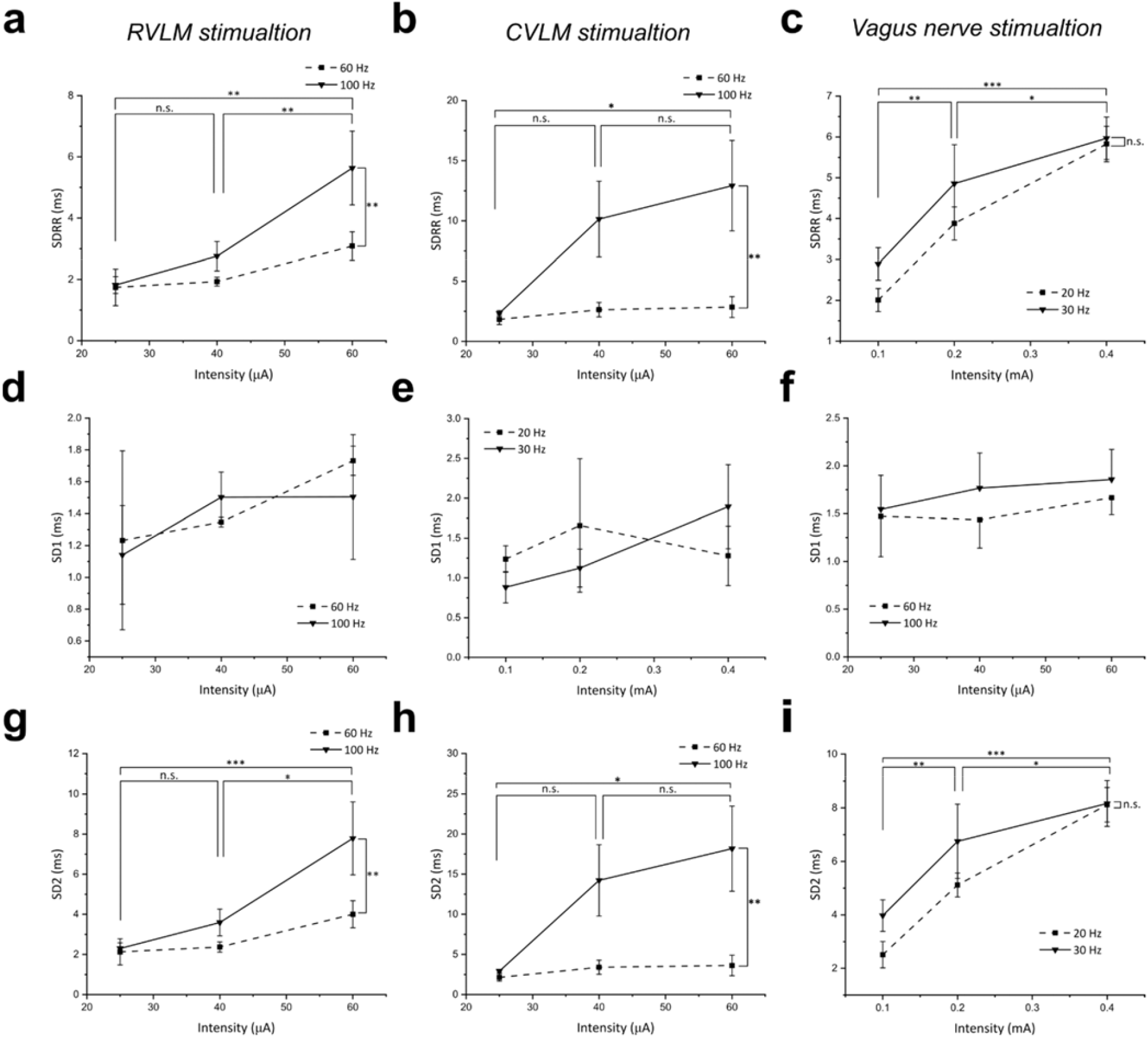
Heart rate variability of open-loop stimulation. Left column, RVLM stimulation; middle column, CVLM stimulation; right column, vagus nerve stimulation. (**a-c**) Standard deviation of RR interval during stimulation. (**d-i**) Quantitative SD1 and SD2 values from Poincaré plots during stimulation. \**P* < 0.05, \*\**P* < 0.01, \*\*\**P* < 0.001, n.s.: not significant by Fisher LSD multiple comparisons test following two-way ANOVA. Data are presented as mean ± SEM.

Since sympathetic nerve trunks innervate the local region of the cardiac ventricles, medullary stimulation may also induce dromotropic effect. Figure 2g-i shows the percentage changes of PR interval from baseline. In RVLM stimulation, the shortened conduction time was enhanced by increased stimulation frequency and intensity. The CVLM stimulation evoked no significant dromotropic changes at all levels of stimulation parameters Vagus nerve stimulation (VNS) evokes cardioinhibitory effects by enhancing parasympathetic nerve activity. To compare this function with CVLMstimulation, the right cervical vagus nerve of rats was stimulated at 20 and 30 Hz and at intensities of 0.1, 0.2, and 0.4 mA. Figure 2c shows the cardiac effects of vagus nerve stimulation at the combinations of every parameter. While the chronotropic response is analogous to the CVLM stimulation, the increasing VNS intensity evoked a more pronounced bradycardic effect at both 20 and 30 Hz (Fig. 2f). Although no significant difference between the two frequencies was found, the curves of average chronotropic responses showed the enhanced tendency for bradycardia at the higher frequency of VNS stimulation. The amplitude (Fig. 3c) and long-term variation (Fig. 3i) of heart rate change have the same characteristics, with the effects saturating at 0.4 mA. Figure 3f demonstrates the same short-term variation at all levels of VNS. For dromotropic effects, the prolonged PR interval effects have a threshold of 0.2 mA, with conduction block enhanced at the higher frequency (Fig. 2i).

### Closed-loop heart rate regulation

Selecting suitable stimulation parameters could provide desired cardiac effect. We were further motivated to test this control ability with simple robust on-off control algorithms in the closed-loop medulla stimulation system (Fig. 1). This on-off controller toggled the stimulus when the heart rate trajectory crossed the target value. Figure 4 shows examples of the dynamic behaviors of controlled heart rate via the RVLM stimulation, CVLM stimulation, and VNS. Note that the target value was set by the peak value of heart rate in open-loop stimulation.

**Figure 4.**
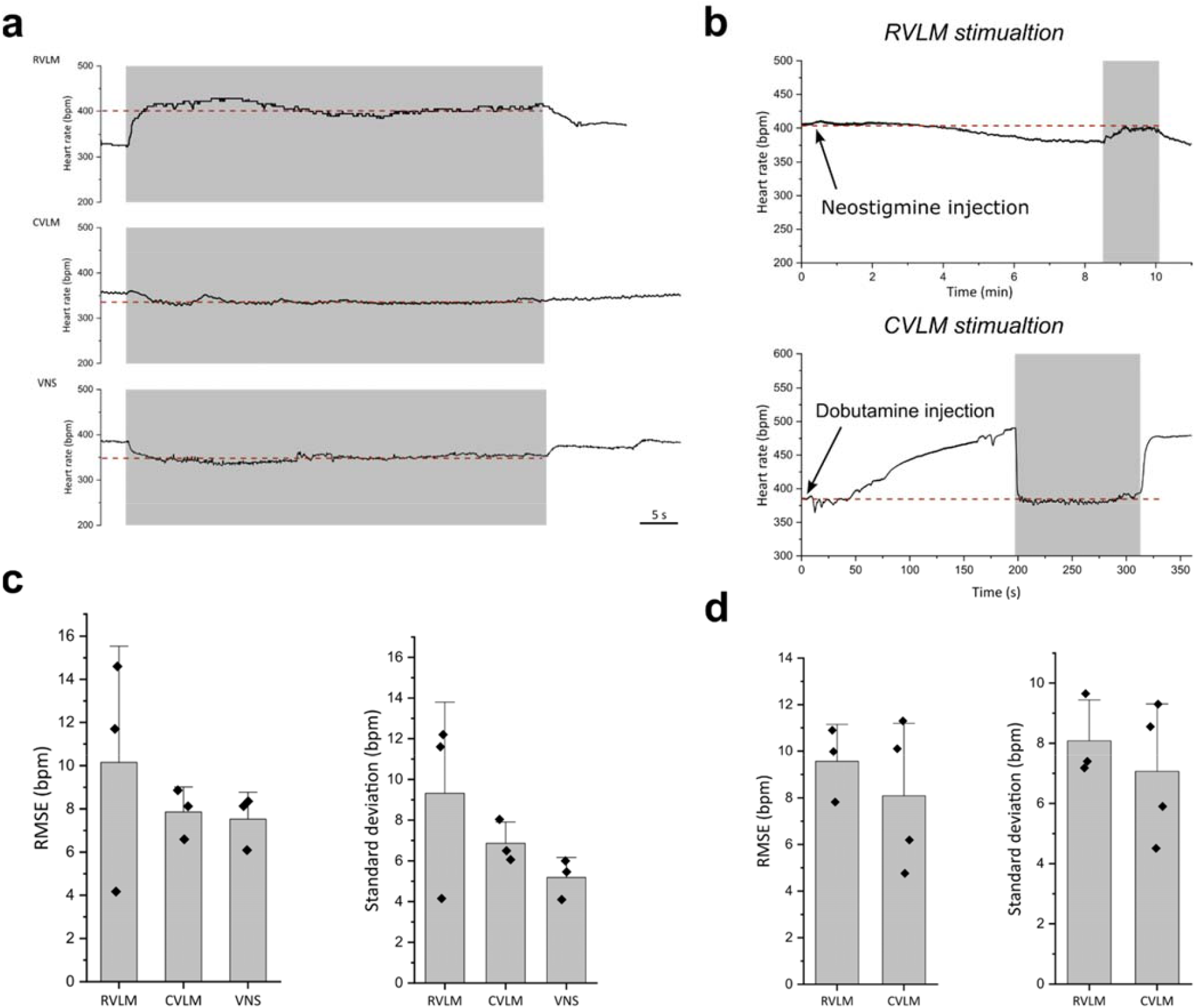
Evaluation of closed-loop heart rate regulation. (**a**) Examples of heart rate control in about 1 min (grey area) at target value of 120% and 90% (dashed line) from baseline via electrical stimulation in RVLM, CVLM and VNS. (**b**) Examples of restoring heart rate to baseline (dashed line) in autonomic imbalance caused by neostigmine and dobutamine injection. During electrical stimulation (grey area), the target values were set as baseline heart rate. (**c, d**) Evaluations of control effect including RMSE and standard deviation. Data are mean ± s.d. with individual animal data (diamond dots).

The control performance was quantified in terms of root mean square error (RMSE) and standard deviation (Fig. 4c). The performance evaluation during the control period was conducted in offline analysis. The RMSE values of RVLM, CVLM, and VNS were 10.2 ± 3.1, 7.9 ± 0.7, and 7.5 ± 0.7 bpm, respectively. Compared to the VNS, closed-loop medulla stimulation modulated the heart rate with higher fluctuations corresponding to standard deviation, although the values of these standard deviations can be negligible due to the high resting sinus rhythm of rodents. Overall, these results demonstrate that precise heart rate control can be achieved in closed-loop stimulation. To determine whether the medulla stimulation can restore autonomic imbalance, the animals were intraperitoneally injected with the acetylcholinesterase inhibitor neostigmine or β_1_ receptor agonist dobutamine to simulate imbalanced autonomic tone. Figure 4b shows a representative heart rate trajectory obtained from these two experiments, demonstrating a reduction of drug effects on heart rate. Note that the stimulus frequency was set according to sustained chronotropic drug effect, ranging from 60 to 100 Hz. As shown in Fig. 4d, the heart rate regulation induced by closed-loop medulla stimulation in drug-administered rats could track the target value of baseline heart rate with slight fluctuation. The results in administered rats were comparable to those in normal rats, demonstrating that the electrical stimulation of the medulla is effective even during imbalanced autonomic activity and can precisely normalize heart rate to its healthy state.

## Discussion

The RVLM region tonically excites preganglionic neurons and maintains the resting vasomotor tone, and also accepts GABA-mediated input from CVLM neurons. This study demonstrates that cardiac responses are primarily driven by the stimulus frequency rather than the intensity, suggesting that tonic activity of neurons in the ventrolateral medulla can be precisely regulated by modulated stimulation, which provides the potential to control the autonomic output of specific physiological index.

Autonomic control of regional cardiac function ultimately depends on the levels of sympathetic and parasympathetic activities and the interactions between them. Many cardiovascular diseases are associated with sympathetic hyperactivity, which induces the remodeling of the cardiomyocyte and the cardiac plexuses that impact cardiac electrical and mechanical functions. As demonstrated herein, the CVLM stimulation evoked the cardioinhibitory effects via inhibiting sympathetic output, which are different from muscarinic blockade effects of VNS with conduction block. This finding suggests that CVLM stimulation can be deployed as an alternative neuromodulation therapy or a tool to explore cardiac remodeling during the progression of cardiovascular diseases.

Chronic tachycardia and hypertension from sympathetic hyperactivity lead to pathophysiological consequences. In animals and patients with hypertension, myocardial and left ventricular remodeling are associated with elevated sympathetic outflow^30–32^. Accordingly, cardiac remodeling develops cardiac arrhythmias and increases the risk of sudden cardiac death^33^. Traditional pharmacological treatments are known to have unavoidable side effects due to complex, sustained pharmacokinetic processes. With the anti-tachycardic effect, VNS has the potential as an alternative therapy to provide cardioprotective effects for drug-resistant patients. Appropriate bradycardia is necessary to prevent hypotension and maintain physiological activities.

The vagus nerve has a mixed fiber population that is classified as A-, B-, and C-fibers in accordance with diameter, myelination, and activation thresholds. Due to the complex fascicular organization and function of the fibers, the stimulation effect depends on various factors such as frequency, current intensity, pulse width, and electrode interface. Previous studies demonstrated many methods for accurate heart rate regulation, including closed-loop control, cardiac-synchronized stimulation, and electrode design^22–24,34–36^. In this study, simple on-off control strategy shows similar cardiac regulation performance with the previous in both medulla stimulation and VNS. To tonically active neurons in the medulla, the efficacy of control could be improved by optimizing the stimulus to match the desired setpoint of heart rate. The methodology of heart rate regulation in medulla stimulation will be easier and more predictable than traditional VNS and pharmaceutical treatments.

For decades, several devices have been developed as bioelectronic medicines using electrical pulses to communicate with the nervous system to improve health as neural prostheses^1^. Most of which act on the baroreflex and vagus nerve, lacking the scenario of sympathetic neuromodulation. Novel neural engineering was developed to miniaturize electrical devices, decipher and modulate neural signal patterns with closed-loop strategies for the detection of dysfunctional activity and achieving therapeutic effects^37^. Although VLM neurons receive inputs including the paraventricular nucleus of the hypothalamus, nucleus of the solitary tract, lateral tegmental field, and A5 region to regulate sympathetic output. In this study, closed-loop stimulation of the medulla can successfully intervene with tonic sympathetic output and alter the heart rate at arbitrary target values depending on modulated stimulus frequency. This regulation capacity in intentional autonomic imbalance conditions implicates the endogenous feedback regulation of sympathetic output may be limited and insufficient, which can be improved with medulla neuromodulation.

Sustained change of cardiovascular homeostasis involves hormones and sympathetic nervous alterations. Brooks and Osborn have suggested a model stating that the raised sympathetic nerve activity is induced by an increased level of circulating angiotensin II^38^, directly modulating blood pressure via activation of angiotensin II type 1 receptor, thus causing vasoconstriction. The elevation of blood pressure activates the baroreflex, which in turn leads to the inhibition of sympathetic efferents and the decrease of blood pressure. However, this reflex in the nucleus of the solitary tract (NTS) can be reset to allow higher levels of blood pressure via the angiotensin II mediated pathway^39^. The mechanism of central sympathetic activity evoked via circulating angiotensin II remains uncertain, probably involving the circumventricular organs which lack a blood-brain barrier and are rich in angiotensin II type 1 receptors. The RVLM neurons regulate renal sympathetic nerve^40,41^, which facilitates renin release from the juxtaglomerular apparatus. It can be assumed that elevated sympathetic activity and angiotensin II level promote each other and are crucial to the development of hypertension, together with a baroreflex resetting in the NTS. Deep-brain stimulation for sympathetic withdrawal has been shown to effectively reduce blood pressure in refractory hypertension^3,42^ and centrally reset the blood pressure reflex^43^. The present chronotropic results suggest decreased sympathetic activity during CVLM stimulation. Future work will be dedicated to the long-term studies of medulla neuromodulation with blood pressure recording.

## Materials and Methods

### SUBJECTS AND SURGERY

25 Wistar male rats (Japan SLC, Kiwa Laboratory Animals) were used in the experiments (300–500 g). All experiments were approved by the Animal Ethics and Administrative Council of Hokkaido University and Kindai University. The rat was initially anesthetized in an induction chamber with 4–5% isoflurane. Once the animal failed to exhibit the toe pinch reflex, the rat was sustained with inhalation anesthesia of 1-2% isoflurane (RWD Life Science) and mounted in a stereotaxic apparatus. The body temperature was maintained at 37°C by a heating pad (TMP-5b, Supertech Instruments). The rat forelimbs were attached to Ag/AgCl electrodes for electrocardiogram (ECG) lead ?. The ECG signals were filtered at 1.5 to 100 Hz and amplified (MEG-5200MG, Miyuki Giken), followed by digitization at 1 kHz through a PowerLab 16/35 data acquisition device (ADInstruments, Australia).

### OPEN-LOOP STIMULATION

To investigate the main factor influencing chronotropic response, various frequencies and currents were applied to the electrical stimulation of the RVLM, the CVLM, and the cervical vagus nerve. For the ventrolateral medulla stimulation, a small craniotomy was performed above the RVLM (from bregma: 12.36 mm posterior, 2.3 mm lateral) or CVLM (from bregma: 13.20 mm posterior, 2.2 mm lateral) on the right side. A coaxial needle electrode (IMB-9004, Inter Medical, Japan) was inserted at the site at a depth of 8.2 mm from the dura. The electrical stimulation was performed via a stimulus generator (STG-4002, Multi Channel Systems) using the following parameters (60 and 100 Hz, 20, 45 and 60 μA, 0.5 ms pulse width, bipolar square pulses), and each stimulation configuration was repeated three times. In the cervical vagus nerve stimulation experiments, the rat was placed in the supine position. After the incision was made at 1 cm lateral to midline, the cervical vagus, jugular vein, and common carotid were exposed by blunt dissection. Following gentle isolation of the right vagus nerve from the carotid sheath, a hook electrode (IMM2-5050, Inter Medical, Japan) was placed around the nerve and secured with drops of liquid paraffin. The anode cephalad to cathode vagus nerve stimulation (VNS) was biphasically delivered at 0.2 ms pulse width to evaluate combinations of frequency and intensity (20 and 30 Hz, 0.1, 0.2 and 0.4 mA) with three repetitions.

### CLOSED-LOOP STIMULATION SYSTEM

A multi-I/O real-time processor (RX8, Tucker-Davis Technologies) loading the custom controller routine detected R wave threshold for computing real-time heart rate and generated the trigger to a stimulus generator (STG-4002, Multi Channel Systems) by an on-off closed-loop law (Fig. 1).

Based on the findings from the pilot open-loop stimulation experiments, the high stimulus frequency of 100 Hz can rapidly induce cardiac effects and saturate heart rate change. To investigate the maximum control ability of heart rate via medulla stimulation, the closed-loop stimulation with 100 Hz stimulus was set to sustain heart rate at desired values of 120% and 90% resting heart rate for tachycardia and bradycardia effect separately. Additionally, the animals were intraperitoneally injected with neostigmine (0.5 mg/kg) or dobutamine (0.4 mg/kg) to explore the control ability in the imbalanced autonomic environment with sympathetic or parasympathetic hyperactivity, respectively. After the heart rate was stably altered, the closed-loop stimulation was performed with the target value of the baseline heart rate. In these conditions, the appropriate stimulus frequency was calculated from the difference between altered heart rate and baseline heart rate to prevent saturation or oscillation in the control system caused by inter-individual variations of drug effect.

### DATA ANALYSIS

For each combination of parameters, ECG signals were continually recorded and divided into 10 s pre-stimulation baseline, and 10 s of cardiac response during the period of stimulation, and post-stimulation. The offline analysis on LabChart software with ECG analysis module was used to detect waves, beats, and calculate intervals. The peak-to-peak RR intervals were used to evaluate the overall chronotropic effect was calculated from the relative change between baseline and period of stimulation. The dromotropic effect was determined via the obtained beat-to-beat PR intervals. To evaluate the efficacy of stimulation, indices of heart rate variability (HRV) in the time domain are presented, including standard deviation of RR intervals (SDRR), and the SD1 and SD2 of the Poincaré plot. The efficacy of heart rate regulation was evaluated by the steady-state error (SSE), defined as the root mean squared error, which was calculated from the target heart rate and the real-time heart rate during closed-loop stimulation. The square root value avoids counteracting errors when oscillations occur around the target value. The standard deviation was calculated to evaluate the stability during the control period. To assess the speed of heart rate response, the time constant was obtained using the non-linear fitting function of the exponential function in OriginPro.

All data are presented as mean ± standard error of the mean. The statistical tests used are explained in the legend of each figure. *P* < 0.05 was considered as statistically significant in all cases. Statistical analyses were performed with OriginPro software.

## Author contributions statement

Y.X. and Y.T. contributed to the conception and design of the study and were involved in the interpretation of the study results. Y.X. was responsible for all stimulation experiments and closed-loop control programming. Y.X. wrote the first draft and illustrated the figures. Y.T., M.C., and M.M. supervised the work and edited the manuscript. All authors contributed to manuscript revision and approved the submitted version.

## Additional information

The authors declare no competing interests.

## Acknowledgements

This work was supported in part by JSPS KAKENHI Grant Numbers JP23K27481 (Y.T.), JP23K18250 (Y.T.), JP24H01999 (Y.T.), JP24K18605 (M.C.), and JP23H04939, AMED under Grant Number JP24gm6510015 (Y.T.) and JP24zf0127004 (Y.T.), Astellas Foundation for Research on Metabolic Disorders, the Japan Epilepsy Research Foundation, Lotte Foundation, the Uehara Memorial Foundation, and Ono Pharmaceutical Foundation for Oncology Immunology and Neurology.

## Notes

### Competing Interest Statement

The authors have declared no competing interest.

